# Bioassay of derived components from venom of Iranian medically important scorpions to identify the bradykinin potentiating factors

**DOI:** 10.1101/210856

**Authors:** Hamid Reza Goudarzi, Ali Nazari, Mojtaba Noofeli, Maedeh Samiani

## Abstract

The venom of animals, including snakes, scorpions and spiders is a complex combination of proteins, peptides, and other biomolecules as well as some minerals. Among the biomolecules, some peptides prevent converting of angiotensin 1 to angiotensin 2 by inhibiting of the Angiotensin Converting Enzyme (ACE) and finally reducing the blood pressure in the victims. The aim of the present study was to isolate venom components of the three species of Iranian medically important scorpions and to study the bradykinin potentiating effect of them. Separation of the venom components for each scorpion was carried out using high performance liquid chromatography (HPLC). The range of fractions (zones) obtained in several replicates on Guinea pig ileum and rat uterus tissues were performed using organ bath instrument. The bioassays were resulted in the peptides including Z_1_ and Z_2_ regions of venom chromatogram of the *Hottentotta sulcyi*, Z_2_ for *Odontobutus doriea* and Z_2_ and Z_3_ in *Mesobutus eupeus* venom demonstrated bradykinin potentiating effect.

## Introduction

There are about 170000 species of venomous animals in the world; the most well-known are snakes, scorpions, spiders and a group of bees. It is estimated that the total venom of these animals contains more than 40,000,000 proteins and peptides [^1^]. Some of the venom compounds are non-enzymatic proteins that bind to specific ion receptors and channels in the injured body and act as an agonist or antagonist which may lead to neurotoxic disorders, cardiotoxicity or tissue necrosis effects [^2^]. Although venoms are recognized as a pathogen in the body, they can also be considered as a healing substance [^3^]. The products derived from creatures have been used since ancient times, so that traditional medicine and ancient medical texts refer to the use of toxins in treatment of various diseases. A classic example of the successful relationship between medicine science and toxinology is the discovery and development of a blood pressure lowering drug. [^4^]. The first time in 1960 a scientist named Ferrera identified and extracted peptides from *Bothros jararaca* snake, which enhanced bradykinin effect and was involved in lowering blood pressure. Further studies on these peptides led to the production of a hypertension drug called captopril. This drug, which is widely used worldwide, reduces blood pressure by inhibiting angiotensin converting enzymes (ACE) [^5^]. Scorpion venom contains a complex combination of mucopolysaccharides, hyaluronidase, phospholipase, serotonin, histamine, inhibitor enzymes, peptides and proteins. These polypeptides are classified into two groups: 1-peptides with Disulfide Bridge and 2-peptides without Disulfide Bridge. The first group is containing peptides that linked about 30-70 amino acids by 3-4 disulfide bonds. This group mainly includes peptides that affect cell membrane channel activity [^6^]. The second group peptides lack the cysteine residue in their sequences and thus lack the disulfide bond. The molecular weight of this group of peptides is in the range of 2500-3000 Daltons. One of the characteristics of this peptide group is the presence of proline residues in their C-terminal end. This group has milder toxic effects than the first group and has antimicrobial, hemolytic and bradykinin potentiating effects [^7^].Studies have shown that the proline amino acid residue in the C-terminal fragment play an important role in enhancing the effect of bradykinin. Thus, recent peptides due to structural and pharmacological properties are considered as one of the most important issues in the biosynthesis of antihypertensive drugs [^8^]. Studies on *Tityus serrulatus* scorpion venom showed that the bradykinin potentiating factors could effect on blood pressure through both inhibition of ACE activity and bradykinin receptor synthesis [^9^]. Based on the reported clinical signs of victims who were stung by scorpion in Iran for the secondary reduction of blood pressure [^10^], the idea of the possible existing of bradykinin potentiating factors in the venom of Iranian medically important scorpion was considered. The current study shows the effect of BPFs isolated from three scorpion species (*Hottentotta sulcyi*, *Odontobutus doriea, Mesobutus eupeus*) based on the bioassay of venom components using isolated tissue contractions in organ bath instrument.

## Materials and methods

### Species of scorpions

The scorpion species studied were: *Odontobuthus doriea, Hottentotta saulcyi, Mesobuthus eupeus* that provided from Razi Vaccine and Serum Research Institute (RVSRI).

### Extraction and clarification of venom

Extraction of the venom was performed by a weakly electroshock (5 volts). The obtained venom was frozen in a 50 °C freezer and then dried by freeze-dryer instrument. In order to separate impurities such as mucoproteins, epithelial cells and possible contaminants, 30 mg of venom was dissolved in 3 ml of distilled water then centrifuged at 4 °C for 30 minutes. The supernatant was later filtered through a 0.45 syringe filter and finally lyophilized.

### Toxicity test (determination of LD_50_)

The serial dilutions of crude venom of each scorpion were selected from doses 100% live to 100% dead with an incremental factor of 1.25 and injected via IV route into Balb/c mice (20 g). The dilution series was prepared in saline solution and injected into 4 groups of Balb/c mice, 20 grams. The results were calculated using Spearman & Karber and Reed & Muench statistical methods [^11,12^].

### SDS-PAGE

In order to determine the molecular weight range of the components of three crude venoms vertical electrophoresis with condition of separating gel 14% and stacking gel 4%, staining with Coomassie Brilliant Blue (G250) was carried out.

### Chromatography RP-HPLC

The High-performance liquid chromatography (HPLC, Amersham Bioscience; ÄKTA prime plus) was used to separate the components of the venom. The chromatographic conditions consisted of columns C18 (Agilent; 4.6 × 250 mm), HPLC grade water (A) and acetonitrile (B) with 0.05% trifluoroacetic acid (TFA) as moving phases at the gradient of 5-80 % (B), and a flow of 1 ml / min. The volume and concentration of injected venom into the device at a time was 2 mg per 200 μl. After obtaining the chromatographic repeatability, the regions (zones) of fractions were determined and collected.

### Dose-Response

Different values of bradykinin from 1-10 ng/ml were applied on the isolated tissue to determine the effective dose to found the mean contraction of the muscle. Then, the effect of bradykinin enhancement was investigated by different fractions obtained from the scorpion venom.

### Preparation of tissues

#### A) Guinea pig ileum

The Guinea pig 200-400 grams which was given 24 hours of starvation, was used. The animal was anesthetized with Ether in a short time then quickly two parts of the terminal ileum were surgically removed. The tissue was rapidly transferred in Tyrode solution containing 1 μg/ml of atropine sulfate. Then, blood vessels and other probable contents were removed from tissue. Organ bath temperature was set at 37 ºC and then the fractions of each scorpion venom were added 10 seconds before adding synthetic bradykinin to the wells. The isolated tissue was washed three times with Tyrode solution after each test.

#### B) Rat uterus

To prepare the rat uterus tissue, 10 ng diacylbestrol was firstly added in 0.5 ml sterile sesame oil and then injected intraperitonealy to the virgin female rat weighed 200-250 g (to ensure that the ovarian tube was empty from ovocytes and tissues, injection of diacylbestrol was performed according to the protocol). After 20 hours, the animal was anesthetized with Ether, in a short time then quickly two parts of the uterus were surgically removed. The tissue was rapidly transferred into De Jalon solution containing 1μg/ml atropine sulfate. After removing of the connective tissue and blood vessels, the tissues were installed in organ bath wells. The other conditions of test were as mentioned above.

## Results

### Toxicity test

The crude venom toxicity (LD50) of each three Iranian medically important scorpions was evaluated and the results are mentioned in Table 1. The mean of the average was obtained from both Spearman & Karber and Reed & Muench methods.

**Table 1:** Toxicity of the crude venom for three scorpion species

| Species | LD50 (µg/mouse) |
| --- | --- |
| *Odontobuthus doriae* | 14 |
| *Mesobuthus eupeus* | 98 |
| *Hottentotta saulcyi* | 75 |

### SDS-PAGE

The electrophoretic pattern of the three crude venoms demonstrates that the molecular weights of the protein and peptide components are ≤ 9 kDa (Fig. 1).

**Fig. 1:**
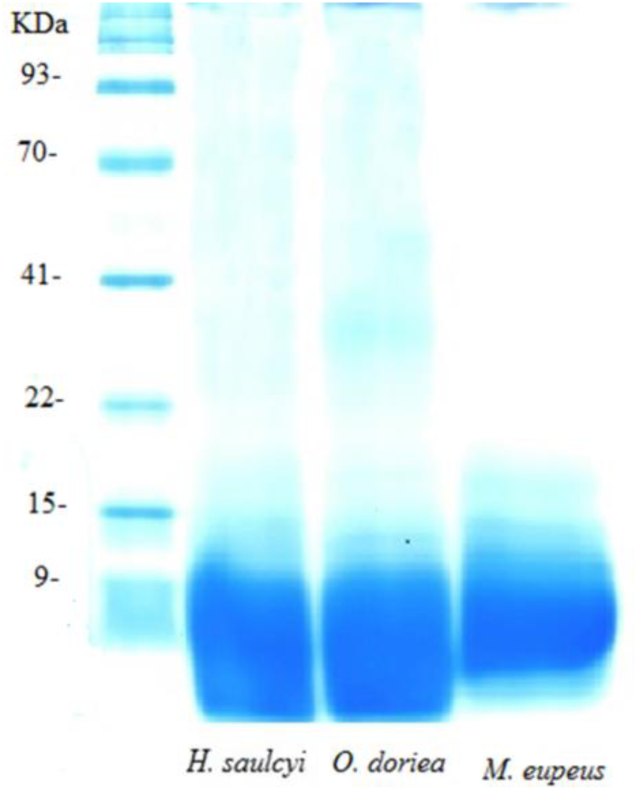
The electrophoretic pattern of crude venoms for scorpions: *Hotenttotta saulcyi, Odontobuthus dorie* and *Mesobuthus eupeus* using 14% separating gel and 4% stacking gel and staining with Coomassie Brilliant Blue G250, Lader; Sinagen co.).

### RP-HPLC

The following chromatograms (Figures 2–4) were repeatedly obtained. Based on these chromatograms, different regions (zones) were isolated and the relevant fractions collected then freeze dried. The regions Z1-Z6 for *H. saulcyi*, Z1-Z4 for *O. doriea* and Z1-Z5 for *M. eupeus* scorpion were selected for bioassays.

**Fig. 2:**
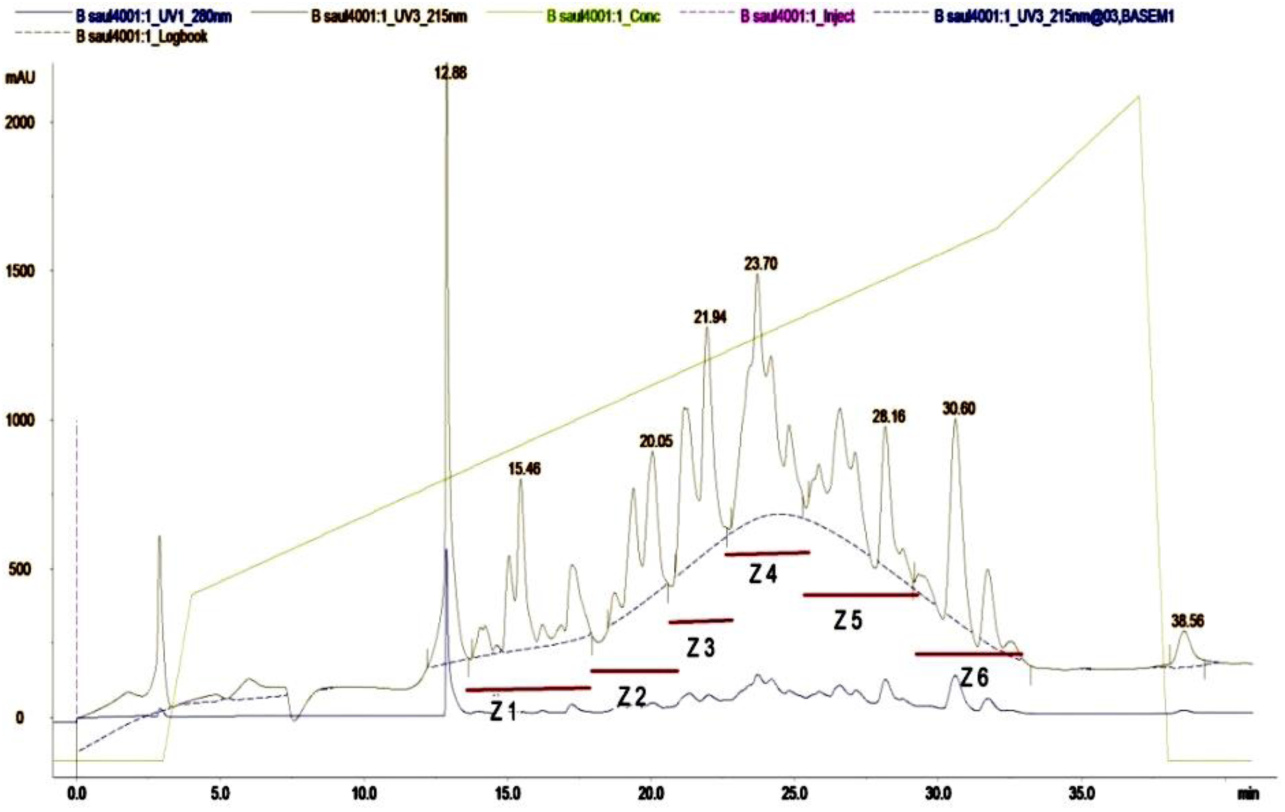
Chromatogram of crude venom fractions of *Hotenttotta saulcyi,* using Reverse-phase HPLC, Amersham Bioscience; ÄKTA prime plus, columns C18 (Agilent; 4.6 × 250 mm)

**Fig. 3:**
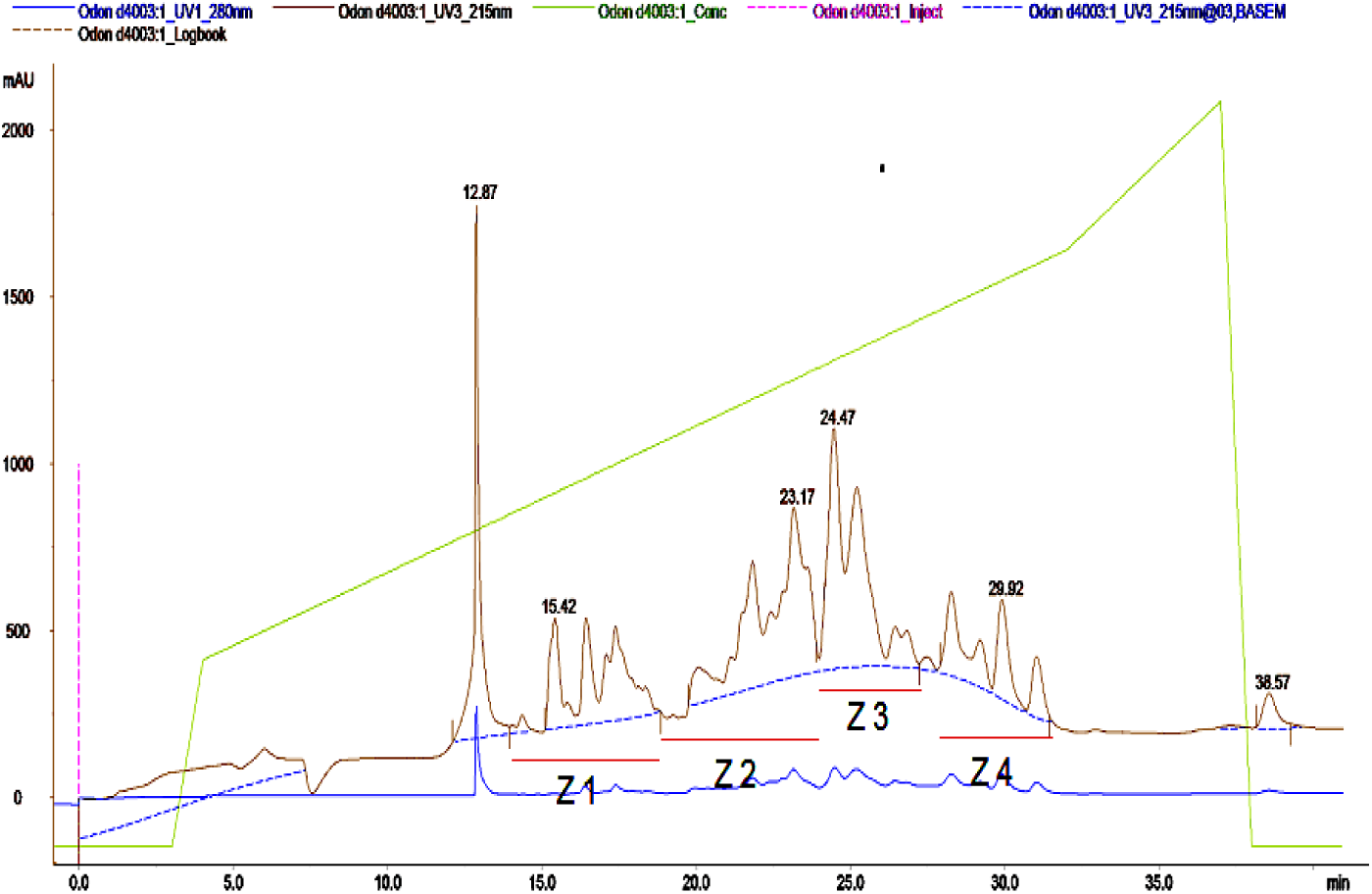
Chromatogram of crude venom fractions of *Odontobuthus doriea*, using Reverse-phase HPLC, Amersham Bioscience; ÄKTA prime plus, columns C18 (Agilent; 4.6 × 250 mm)

**Fig. 4:**
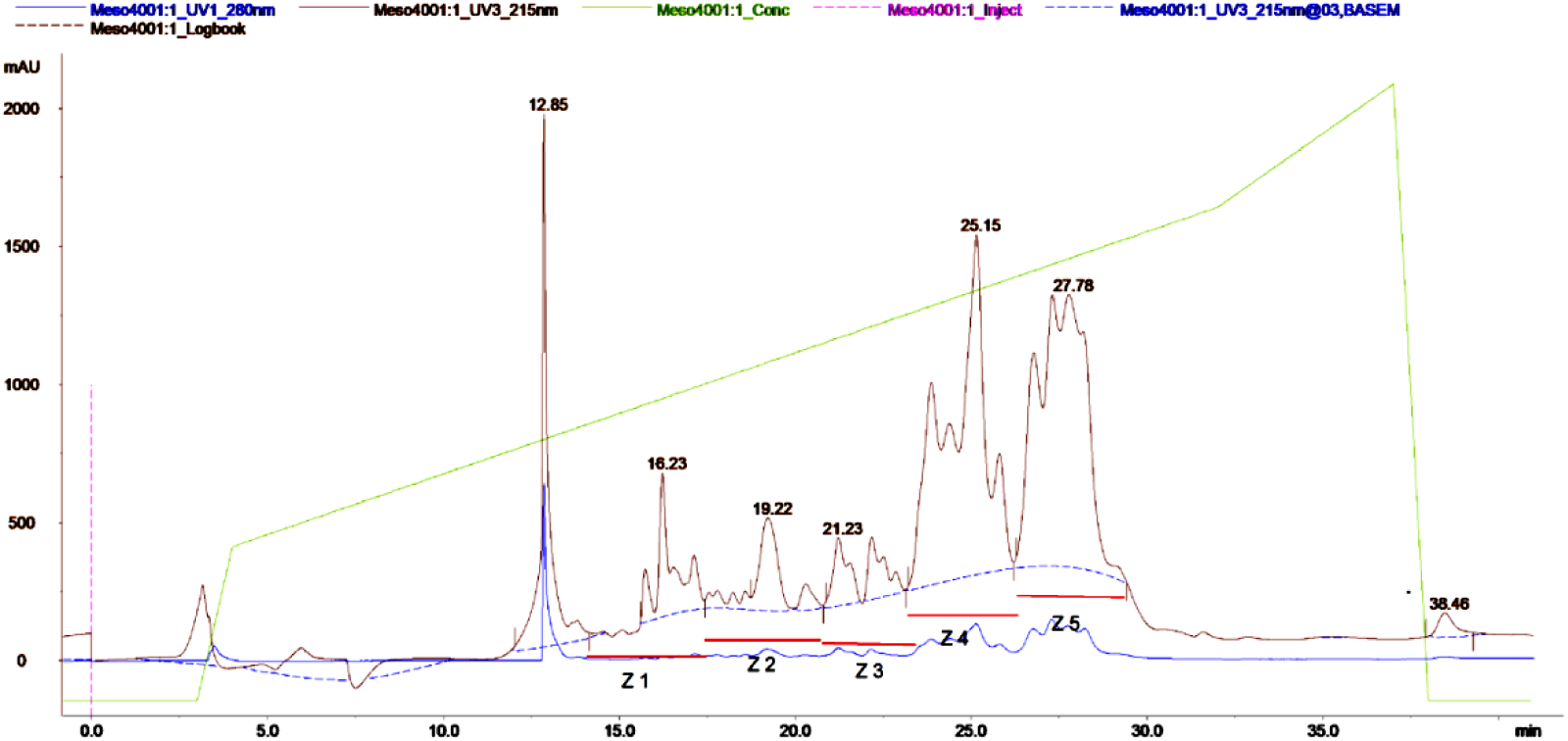
Chromatogram of crude venom fractions of *Mesobuthus eupeus*, using Reverse-phase HPLC, Amersham Bioscience; ÄKTA prime plus, columns C18 (Agilent; 4.6 × 250 mm)

### The zones including toxic fractions

The venom of the Buthidae family species which was studied is strongly neurotoxic. These toxins are almost bounded to acetylcholine receptors. The attached toxin prevents the transmission of neural impediments during neuromuscular involvement and results in sequential contractions. The areas of Z3 and Z4 in the venom of *H. saulcyi*, Z3 of *O. doriea*, and Z4 and Z5 regions in *M. eupeus* illustrate toxic effects. However, in future studies, the aforementioned zones can be purified and each fraction analyzed separately.

**Fig. 5:**
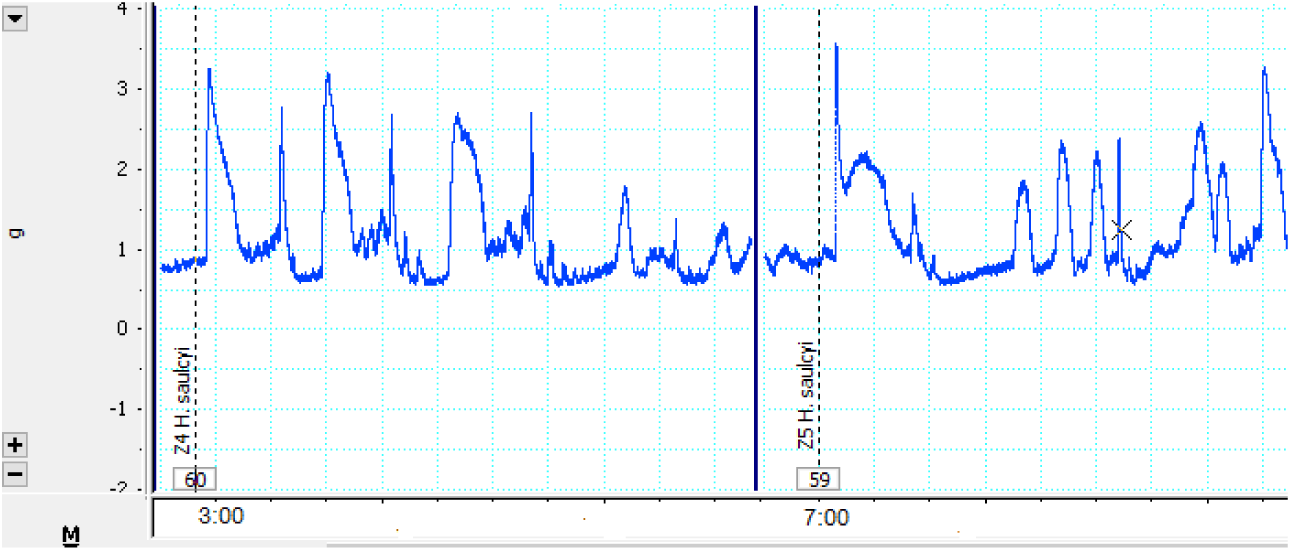
The effect of toxic fractions on isolated tissue which cause severe oscillations in muscle contraction.

### Dose response test results

The results of the dose-response study are shown in Fig. 6. The dose threshold was 1ng/ml with the first dose through a change in muscle tension. However, for bioassay experiment it should be required to obtain a mean dose. In this assay, the midpoint of the muscle stretching on a curve was determined as 4ng/ml which produced a tensile of 2.3 ± 0.1 g.

**Fig. 6:**
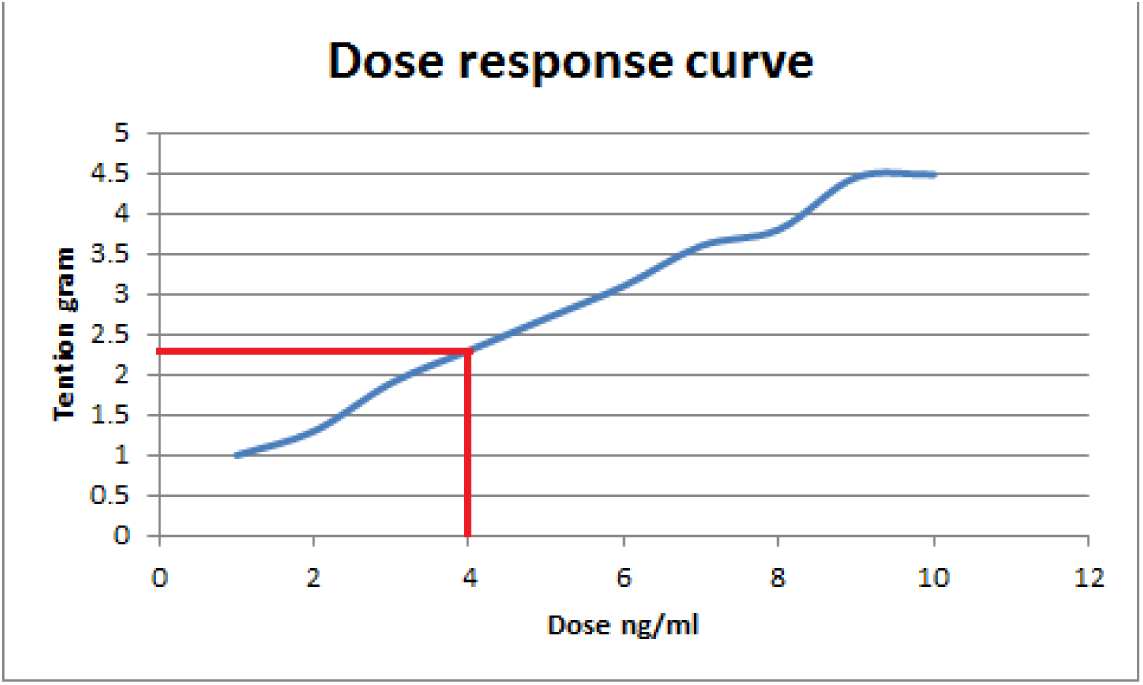
Dose response curve of bradykinin acetate on isolated tissue of Guinea pig ileum

### Bioassays (bradykinin potentiating effects)

The obtained zones including nontoxic fractions were tested on Guinea pig ileum and rat uterus tissues in six replicates using organ bath instrument. The Z1 and Z2 regions of *H. saulcyi* venom fractions, Z2 region of *O. doriea* and Z2 and Z3 fractions of *M. eupeus* venom showed obviously the bradykinin potentiating effect, based on measuring the contraction of isolated tissues.

As shown in figures 7 and 8, respectively, contraction graphs of isolated ileum and uterus tissues, initially added venom fractions did not show any change in the amount of muscle tension, but after 10 seconds when bradykinin was added to the wells (4ng/ml), the amplification pattern was visible compared to only adding venom fractions. The positive amounts of potentiation unit (PU) (the ratio of the increasing in tension amount to the tension rate resulting from the dose-response) in the Table 2 demonstrates significantly existing bradykinin potentiating factors in the venoms of scorpions studied. The potentiating property of venom fractions occurred on both types of isolated tissues, though amounts of PU in Guinea pig ileum was bigger than rat uterus. However, this assumption should be more studied in future.

**Fig. 7:**
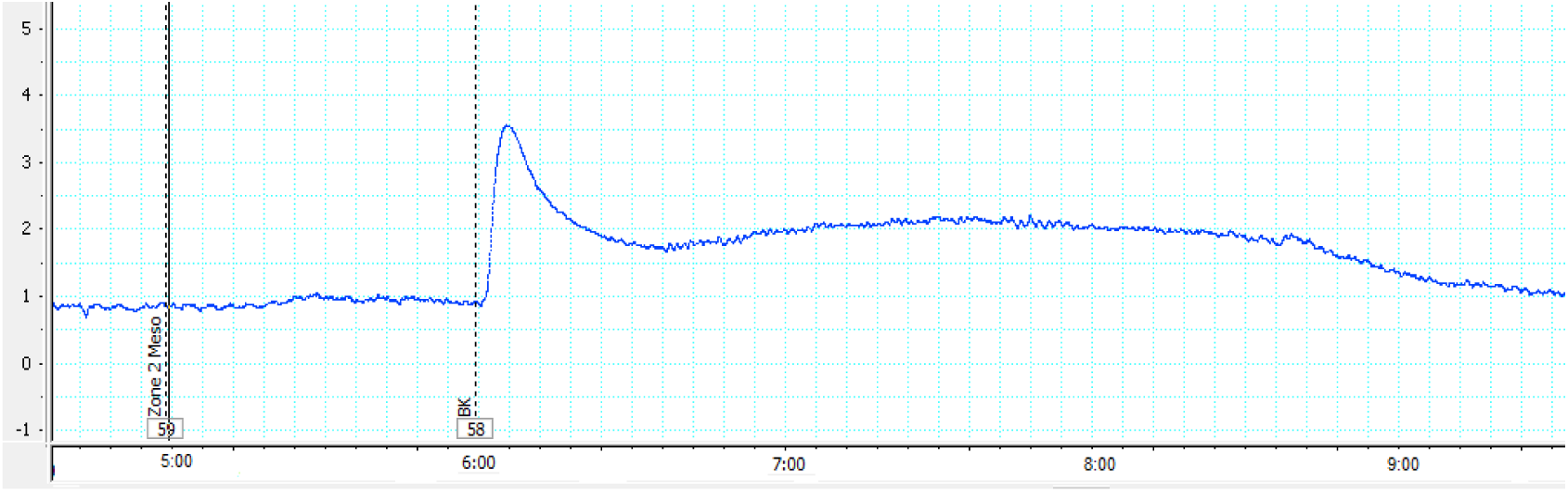
The pattern of the contraction curve for the isolated Guinea pig ileum. It was initially added by venom fractions of *Mesobutos eupeus* scorpion. After 10 seconds, addition of the bradykinin (mean dose) increased the contraction to a maximum of 3.6 g. The time to reach the balance lasted for 35 seconds.

**Fig. 8:**
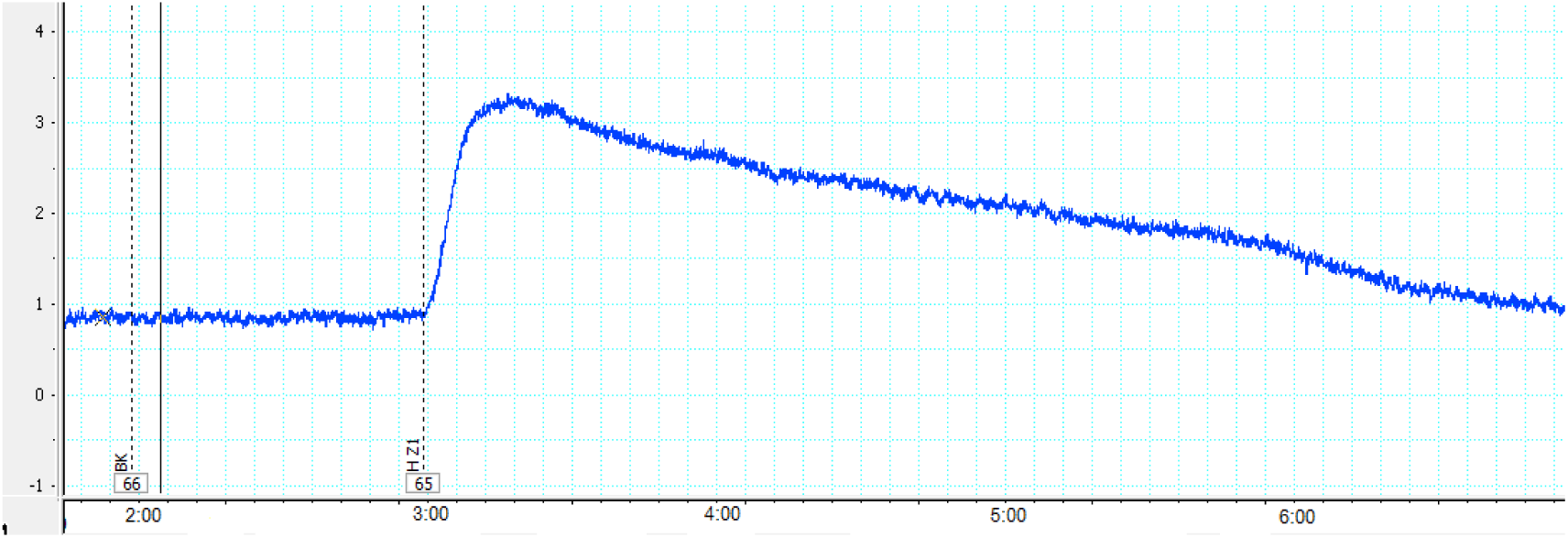
The pattern of the contraction curve for the isolated rat uterus. It was initially added by venom fractions of *Mesobutos eupeus* scorpion. After 10 seconds, addition of the bradykinin (mean dose) increased the contraction to a maximum of 3.3 g. The time to reach the balance lasted for 35 seconds.

**Fig. 9:**
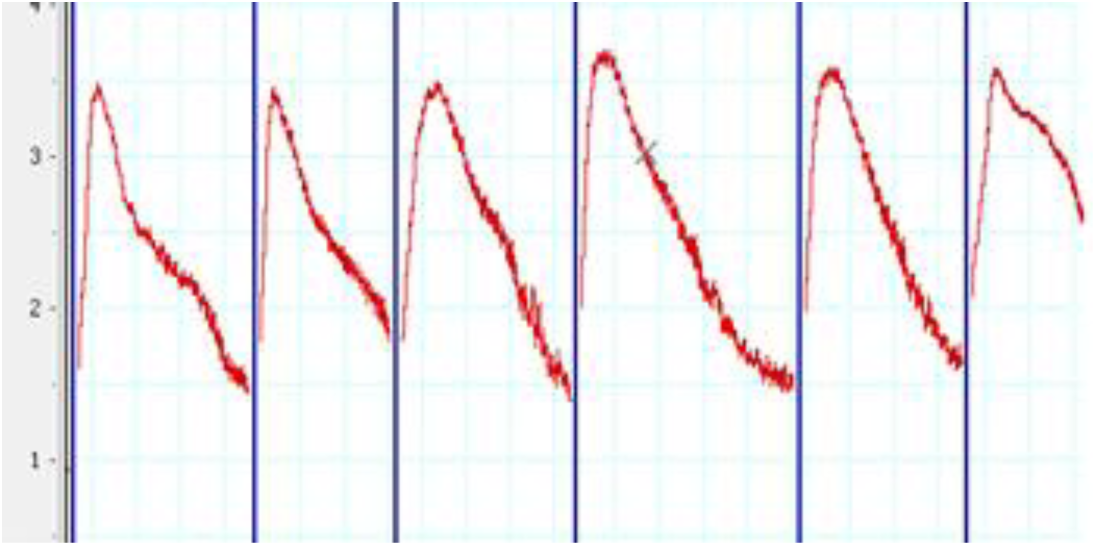
The contraction graphs, potentiating effect of venom fractions on isolated Guinea pig ileum.

**Fig. 10:**
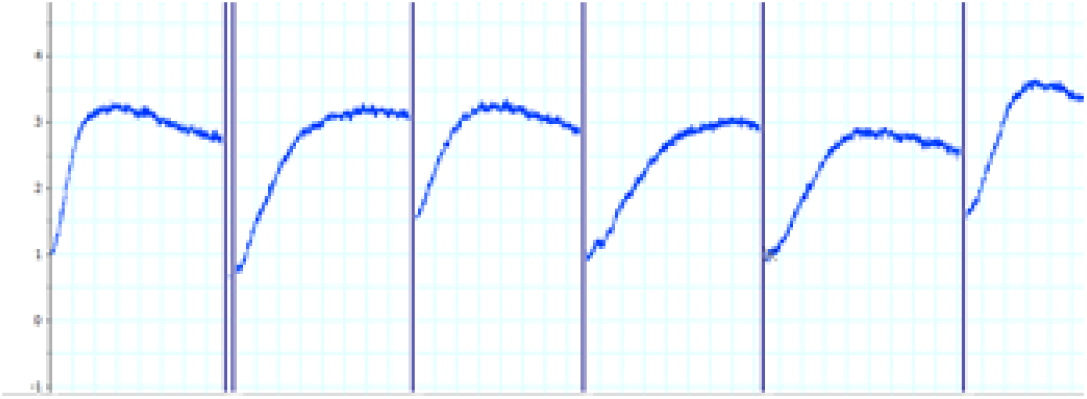
The contraction graphs, potentiating effect of venom fractions on isolated rat uterus.

**Table 2:** Average of potentiation unit, resulted by venom fractions of each scorpion

| Scorpion | Fraction Zones | PU on Guinea pig Ileum | PU on rat uterus |
| --- | --- | --- | --- |
| *H. saulcyi* | Z<sub>1</sub> | 0.26±0.029 | 0.22±0.024 |
|  | Z<sub>2</sub> | 0.44±0.039 | 0.35±0.035 |
| *O. doriea* | Z<sub>1</sub> | 0.52±0.045 | 0.30±0.032 |
| *M. eupeus* | Z<sub>1</sub> | 0.57±0.047 | 0.39±0.037 |
|  | Z<sub>3</sub> | 0.61±0.039 | 0.48±0.032 |

## Discussion

During the last decades, research on venom of scorpions has attracted the attention of toxicologists. Hence, increasing knowledge in this field has been remarkable over the years. The intriguing achievements resulting from the beneficial efforts of researchers in this field are simultaneously known as applied and theoretical tendencies. In addition to isolating and studying the structure of toxic agents involved in humans and animals, also studies have focused on the pharmacological effects of peptides in scorpion venom. It is worth noting that research on low molecular weight peptides and biological activity in snake venom began about three decades before such studies on the scorpion venom, especially on peptides having bradykinin-potentiating activity.

Because of very divergence of the first and second structure of the non-disulfide-bridged peptides (NDBPs) in the scorpion venom, their classification is very difficult based on the amino acid sequences or structural similarity. Therefore, in the relevant sources, the classification based on comparing pharmacological activity, peptide length and building similarity are carried out. Zhian Chan *et al.* (2005) classified these peptides into peptides without disulfide bridges separated from scorpion venom. Although, the same work was previously performed to unify the nomenclature of short chain peptides by Titgat *et al* [^13^].

In recent decade, studies on the bradykinin potentiating-factors from scorpion venom were developed. Ferrera *et al.* (1993) isolated the peptide T as a novel BPP from *Tityus serrulatus* scorpion venom [^14^]. Also, Meki, Nassar and Rochat (1995) separated peptide K12, a bradykinin-potentiating peptide from an Egyptian scorpion venom named *Buthus occitanus* [^15^]. Zeng XC., *et al*. (2000) in a more advanced research, cloned and characterized a novel cDNA sequence encoding the precursor of a venom peptide (BmKbpp) related to a bradykinin-potentiating peptide from Chinese scorpion named *Buthus martensii* [^16^]. In another research by Sosnina *et al.* on venom of a spider species, the near member of arachnid class to scorpions, effect of BPPs from the *Latrodectus tredecimguttatus* in inhibition of carboxycathepsin and its role in preparation of black widow spider venom kininase were studied [^17^].

Surprisingly, in recent years, studies on various pharmacological aspects of isolated BPPs from scorpion venom have been improved. For instances hepato and nephroprotective effects of bradykinin-potentiating factors [^18^], effect of a single dose of a BPF on total protein and albumin in serum [^19^], prevention of hepatic and renal toxicity with [^20^] as well as protective effect of BPF against kidney damage in laboratory animals [^21^].

In the present study, the venoms of three species of Iranian scorpions named *Mesobutus eupeus*, *Hotenttotta sulceyi*, *Odontobutus dorea* were studied for detection and measurement of bradykinin-potentiating activity of their fractions. The mentioned scorpion specimens are native species in Iran; so far none of these scorpion venoms has been investigated by Iranian or other researchers. As a result, the Iranian scorpions are valuable sources to isolate and study on the biological factors along with various pharmacological effects such as bradykinin potentiating peptides. Probably even in the zones with toxic effects, peptides with bradykinin potentiating activity are present which dominate by toxic elements. So, further studies in the future are necessary for the separation and purification of target peptides and then, molecular studies and amino acids analysis.

## Acknowledgment

This study was supported by a grant from the Razi Vaccine & Serum Research Institute, Karaj, Iran. In particular, the authors thank the staff in the Department of Venomous Animals and antisera production.

